# Contrasting roles of host species in tick-pathogen interactions and their influence on Lyme borreliosis hazard across boreal island communities

**DOI:** 10.1101/2025.08.21.671197

**Authors:** Yingying X.G. Wang, Nosheen Kiran, Ilze Brila, Jenni Kesäniemi, Saana Sipari, Eugene Tukalenko, Saija Vuorenmaa, Erin Welsh, Andy Fenton, Tapio Mappes, Eva R. Kallio

## Abstract

Predicting the hazard of Lyme borreliosis (LB) is complex due to host-specific effects on tick and pathogen dynamics. To unpick the contributions of cervids and rodents on ticks and *Borrelia burgdorferi* infection in ticks in boreal region, we analysed data from 41 islands in Finland (2017-2021) with considerable natural variation in the abundances of cervids and rodents, and densities of nymphs (DON: 0-82.5 nymphs/100 m^2^), nymph infection prevalences (NIP: 0-60%) and densities of infected nymphs (DIN: 0-16.5/100 m^2^). Using structural equation modelling (SEM), we disentangled direct and indirect effects of host abundances on LB hazard (i.e., DIN). Rodent abundance was associated with higher DIN through its positive effect on NIP (pathogen amplification), while showing no effect on DON. Cervids increased DON (vector amplification) but decreased NIP (pathogen dilution). Our study suggests that in boreal forests the vectors may be supported by very low cervid densities and, as a virtual absence of rodents did not eliminate the pathogen, the pathogen is likely to persist through very low rodent densities. Our study highlights how islands function as a natural mesocosm for rigorous field studies of complex ecological interactions and SEM can serve as a powerful tool to reveal mechanisms driving LB hazard.

## Introduction

Predicting and managing the hazards caused by zoonotic tick-borne pathogens (TBP) is challenging due to the complexity of TBP circulation. Typically, TBP circulation involves the pathogen, the tick, and multiple host species that may differ in their relationships with both the vector and the pathogen (1,2). In northern ecosystems, the most common tick-borne disease is Lyme borreliosis (LB) (3). LB is caused by the bacterium *Borrelia burgdorferi* sensu lato (s. l.), transmitted by *Ixodes* ticks, of which *Ixodes ricinus* is the most important vector in Europe (2,4). Despite the extensive efforts to quantify the drivers of LB hazard and risk in different ecosystems, the debate over the roles of different types of hosts continues (5–10).

*Ixodes* ticks and TBPs are differently associated with vertebrate host species. The sheep tick (*I. ricinus*) is a multi-host vector that utilises a wide range of host species for blood meals needed for development (larvae, nymphs) and reproduction (adult females) (11). However, the most important host groups differ for each life stage. Larvae feed mostly on small vertebrates, such as rodents and birds, but can also utilise larger mammals (8,12,13). Nymphs utilise a wide range of hosts, whereas adult females require a blood meal from a sufficiently large mammal such as cervids (deer and moose; (12-14)). Meanwhile, the pathogen *Borrelia burgdorferi* s. l. is dependent upon its vertebrate hosts, without whom pathogen circulation is not possible. In particular, small mammals such as rodents are important transmission hosts for *B. afzelii* and *B. burgdorferi* sensu stricto (15,16), the most common LB-causing pathogens in Europe and US, respectively (2,4,17). *Ixodes ricinus* larvae acquire *B. afzelii* from infected small mammals and transmit the pathogen onwards, e.g. to susceptible hosts and humans, typically at nymphal stage. Importantly, deer, which support tick reproduction, are not competent transmission hosts for *B. burgdorferi* s. l. (2,15). Consequently, the relative densities of the different types of hosts, particularly cervids (tick reproduction hosts) and rodents (pathogen transmission hosts), are crucial drivers of vector and pathogen dynamics.

Variation in rodent population density has been shown to translate into variation in pathogen prevalence in ticks, with positive associations between nymphal infection prevalence (NIP) or density of infected nymphs (DIN) and the previous year’s rodent abundance (e.g. 9,10,18,19). Furthermore, as rodents are also important feeding hosts for larval ticks, the rodent density one year has been shown to translate into the following year’s density of nymphs (DON) (10). These findings are largely based on the studies in the temperate zone in Central Europe and North America, where rodent population dynamics are strongly affected by the masting of deciduous forests, which provides food for rodents and supports their over-winter survival, resulting in very high rodent densities (10). In boreal zone in Fennoscandia, however, rodent population dynamics are characterised by multiannual (3-5 years) cyclic density fluctuations, which are mostly driven by specialist predators (20,21). These population fluctuations are synchronous for most small mammals (voles and shrews) and over large areas (21). Typical for these cycles is a very low-density phase (< 1 individual/ha) over an extended time period (over a year) (21). Low densities of transmission hosts may be detrimental for the persistence of the pathogen (22). Hence, such prolonged low-density phases of the cycles may cause the pathogen to be eliminated from the system if transmission from nymphs to larvae via rodents is prevented. Yet, how tick density and infection prevalence in ticks are affected by the rodent population dynamics in Fennoscandia remains largely unresolved.

Tick abundance is often associated with the abundance of large mammals, especially deer, which provide blood meals for adult females, thus supporting tick reproduction. For instance, roe deer (*Capreolus capreolus*) and white-tailed deer (*Odocoileus virginianus*) are regarded as the most important tick-reproduction host in Europe and the US, respectively (23). However, reported associations between deer and tick abundances have been inconsistent. As summarised by (24,25), observational studies have typically shown a positive association between tick abundance and deer (26), although deer density is not necessarily (linearly) translated into the density of nymphs (DON) (24,27). Moreover, experimental studies that reduce deer abundances have provided especially variable results, ranging from an increase to a drastic decrease in tick densities (28-30). Consequently, the role of deer in determining tick abundance is still considered one of the key knowledge gaps in understanding the ecology of Lyme borreliosis (23). Importantly, deer provide blood meals not only to adults but also to other life stages of ticks (11). Since deer are not suitable transmission hosts for *B. burgdorferi s*.*l*. (2,15), their presence potentially diverts immature ticks from feeding on infectious hosts, disrupting transmission. Consequently, a high density of deer relative to the density of rodents may lead to a decrease (i.e., dilution) of pathogen prevalence in ticks (1,6,7). Quantifying the dynamic interplay between different types of hosts, ticks, and their pathogens remains vital for understanding the effects of changing host community composition on the zoonotic hazards of TBPs.

Our aim was to quantify the effect of different host types, each with a different role in supporting the tick and the pathogen, on the hazard of Lyme borreliosis in a boreal ecosystem in Fennoscandia. We collected data on the abundances of rodents and cervids, and the abundance of ticks, and the prevalence of *B. afzelii* in ticks, using 41 islands in Finland as a natural mesocosm (Map S1 in Supplementary information). The relatively close proximity of the islands means any potential effects of environmental variation on ticks are reduced, as climatic conditions and forest habitats, dominated by pine, spruce, and birch, are similar across the islands. However, each island was expected to represent its own multi-host-tick-*Borrelia* system, as the movement of the hosts, ticks, and pathogen between the islands is hindered by water. Hence, tick and pathogen abundances on each island are likely to reflect transmission processes determined by local host densities on each of those islands. The natural variation in the densities of the two types of hosts, ticks and pathogens, across the islands provides an ideal opportunity to disentangle the contributions of each host type to tick and pathogen transmission across the system.

The interactions between the hosts, tick, and the pathogen in this system may comprise a network of interdependent processes, where direct effects cascade through indirect pathways. To address these complexities, we employed a Structural Equation Modelling (SEM) framework, which allows simultaneous quantification of both direct and indirect effects (31,32), to quantify the drivers of Lyme borreliosis hazard. Specifically, we examined whether the local abundances of both host types (cervids and rodents) are associated with the local density of nymphs (DON) and nymph infection prevalence (NIP), which together determine the density of infected nymphs (DIN). We hypothesised that (i) DON is positively associated with the preceding year’s abundance of tick-reproductive hosts, i.e., cervids (7,26), and that (ii) DON is also positively associated with the abundance of rodents, which serve as key hosts for larval ticks (9,10). Additionally, we hypothesised that (iii) NIP increases with increasing rodent abundance due to their role as competent pathogen reservoirs (i.e. pathogen amplification) (9,10,18,19), and (iv) NIP decreases with cervid abundance, consistent with a pathogen dilution effect from non-competent hosts (1,6,7). Consequently, we expect that cervid abundance would have countervailing indirect effects on the density of infected nymphs (DIN), with a positive indirect effect via DON, and a negative indirect effect via NIP.

## Material and methods

### Study sites and study design

The study was conducted on 41 islands (55 sites) in the Porvoo archipelago in Southern Finland during 2017-2021. The sizes of the islands varied between 0.5 and 966 ha. All study islands were located within an area of ca. 50 km^2^ (Map S1, Supplementary Information). In small islands (<4 ha), the sampling covered most of the island, whereas in larger islands, each study site covered approximately a 3-4 ha area within which all the data were collected. In most study islands there was only one study site, but in some larger islands up to five sites were examined (Map S1), with an inter-site distance of at least 500 m.

The study design (Figure 1) encompasses the life cycle of the ticks in Finland (33,34), with adult ticks seeking a host from early spring in one year (Year_t_) and producing eggs, which hatch to larvae that seek for host in the summer-autumn of that year (Year_t_). If successful, the larvae moult to nymphs that quest for a host from the spring of the next year (Year_t+1_). The data collections included (i) rodent trappings in late August – mid-September (hereon “autumn”) in 2017-2020 (Year_t_), (ii) tick collections in the following May (hereon “spring) in 2018-2021 (Year_t+1_), and (iii) large herbivore dung surveys in spring (Year_t_) and in autumn (Year_t_) together with the tick and rodent data collections in 2019-2021 (Figure 1). The rodent trappings were carried out in 24 islands (32 sites) in autumn 2017, 29 islands (36 sites) in autumn 2018, 29 islands (34 sites) in autumn 2019, and 23 islands (26 sites) in autumn 2020. Subsequently, the same islands and sites were visited in the following spring for tick collections (Figure 1). The numbers of sites/islands visited varied between years to maximise the number of islands examined during the study with the available resources and permits from landowners.

**Figure 1.**
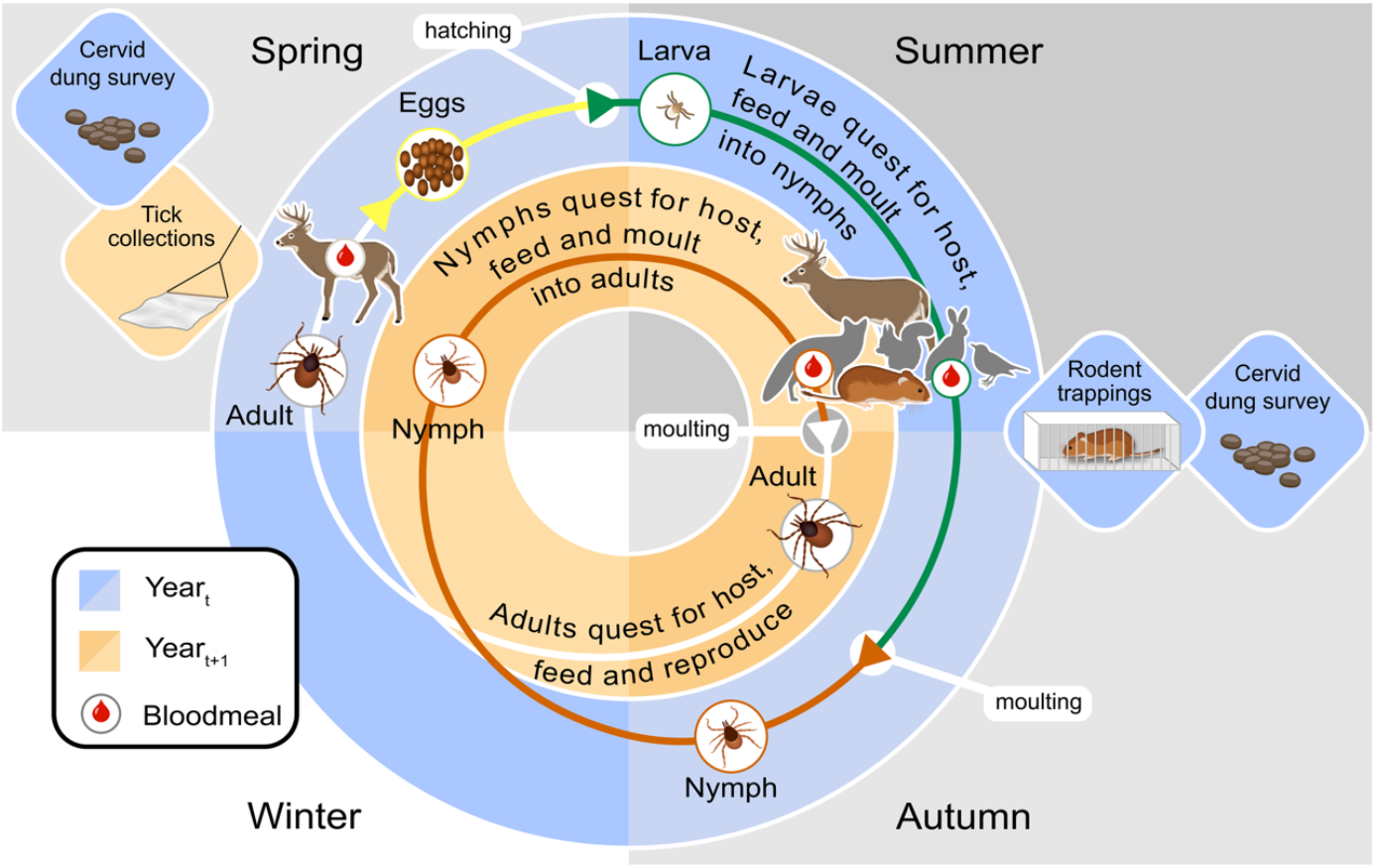
Schematic figure of the study design together with the tick life cycle in the study region. Tick larvae quest for a host in summer-autumn, and in the following spring the tick quests for hosts as a nymph. Consequently, in autumn (late August-September) rodent trappings (2017-2020) and dung surveys (2019-2020) were carried out, whereas in spring (mid-late May) ticks (2018-2021) were collected and dung surveys (2019-2021) carried out. Figure created by MārisGrunskis/@PHOTOGRUNSKIS.

### Data collection

#### Rodent trapping and sampling

Rodents were trapped each autumn in 2017-2020 using Ugglan special live traps (Grahnab, Sweden) that were supplemented with snap traps in large islands. Depending on the size of the island, a total of 8 to 45 traps were located with ca. 10-20 m intervals along with two or three transects. The traps were set in the afternoon/evening and checked and removed on the following day. In the live traps, sunflower seeds (*Helianthus annuus*) were used as bait and food, while a piece of potato was used to provide water. In the snap traps, a piece of rye bread was used as bait. Live-trapped individuals were handled either on the island of the capture or they were transferred into a sampling laboratory for handling. Individuals that were snap-trapped were recorded and individually stored at -20°C. The species of each captured rodent was identified to species. Of the 1195 rodents captured, 273 were caught with snap traps.

#### Cervid dung surveys

The most common hosts supporting tick reproduction in the study area are three species of cervids: the white-tailed deer (*Odocoileus virginianus*, which was introduced in Finland in 1920), the roe deer (*Capreolus capreolus*) and the moose (*Alces alces*). To estimate the abundance of cervids, we conducted dung surveys from spring 2019 until autumn 2021. Dung piles were counted on a 1-meter-wide square-shaped track, with each side approximately 20 meters, in total 80 meters. Depending on the size of the island, 2-4 sampling squares were fitted per site. Due to uncertainties in separating roe deer and white-tailed deer dung and the expected ecological similarities of deer and moose, the cervid dung pile counts were summed and used as proxy for total cervid abundance on the site.

Other mammalian species that may host ticks in the study systems include the raccoon dog (*Nyctereutes procyonoides*), red fox (*Vulpes vulpes*), domestic dog (*Canis familiaris*) and two species of hare (*Lepus sp*.). However, the dung piles of these species were observed with very low frequencies or on only a few islands, and thus these species were not considered in this study.

#### Tick collection

Ticks were collected in May of 2018-2021 in the same locations that rodents were trapped in the previous autumn. Ticks were collected using the drag flagging method, where a 1 m^2^ white flannel is dragged on the vegetation. The flag was checked every 10 meters, ticks were counted per life stage (nymphs, adult females, and males), marked, and removed with tweezers, and placed in microtubes with 70% ethanol. Large numbers of larval ticks were not counted individually but approximated, and they were removed from the flag with masking tapes and placed in sealed plastic bags. The flag dragging was carried out in total for 200-400 m per site, depending on the abundance of ticks: if less than 40 nymphs/200 m were collected, the dragging continued up to 400 m. In very small islands, smaller numbers of drags were done and, in some sites, where a low number of ticks was collected, extra drags were done. The ticks were stored at -20°C in ethanol, until individuals were sorted according to life stage and identified to species level based on morphological characteristics under a stereomicroscope using identification keys (35,36). Nymphs were used for pathogen screening (see below).

### Ethics statement

All procedures for mammals followed the Finnish Act on the Use of Animals for Experimental Purposes, approved by the Finnish Animal Experiment Board (permission numbers ESAVI/7256/04.10.07/2014 and ESAVI-3981-2018). According to the Finnish Act on the Use of Animals for Experimental Purposes (62/2006) and a further decision by the Finnish Animal Experiment Board (16th May 2007), traps that instantly kill the animal, is not considered an animal experiment and therefore requires no animal ethics license from the Finnish Animal Experiment Board. Tick collections do not require any ethical permissions. All data collection took place with the permission from landowners.

### Pathogen and tick species detection from nymphs

Individual nymphs were used to extract total DNA using the ammonium hydroxide (NH_4_OH) (Sigma Aldrich, Munich, Germany) method (37) described in the supplementary methods (Supplementary information). A negative control (extraction solution without a nymph) was included after every five samples. The lysate was stored at -20°C until further use.

Nymphs were first screened for *B. burgdorferi* s. l. using a generic *Borrelia* qPCR assay targeting flagellin encoding region (38). Nymphs positive in the first assay were examined also using a *Borrelia afzelii-*specific qPCR assay also targeting flagellin gene (39). A subset of 350 nymphs was identified for tick species using a species-specific duplex real-time quantitative PCR (qPCR) assay (40). All used primers and probes are listed in Table S1 and the details of the qPCRs are provided in the Supplementary Information (methods in SI).

The number of nymphs examined per site per year varied due to the variation in the numbers of ticks collected. When the number of nymphs was high, we randomly selected up to *ca*. 40 individuals per site per year for pathogen detection. If less than five nymphs were collected from a site in a year, the pathogen screening was not carried out for those ticks to avoid problems with inferring pathogen variables based on very small sample numbers.

### Variables in statistical analyses

For the statistical analyses the following metrics per site per study year were calculated. *Rodent abundance* (a proxy for the rodent density) per site per autumn was calculated as the number of captured rodents (including all three species) divided by the number of traps used, multiplied by 100 (=abundance/100 trap nights). *Cervid dung abundance* (a proxy for the abundance of cervids) was calculated as the number of dung piles observed divided by the observation distance, multiplied by 100 m (=dung/100 m). The density of nymphs per 100 m^2^ (*DON*) was calculated based on the number of nymphs collected, divided by the dragging distance per site per year and multiplied by 100. Nymph infection prevalence (*NIP*) was calculated based on the number of *B. afzelii* -infected nymphs divided by the number of tested nymphs. The density of infected nymphs per 100m^2^ (*DIN*) was calculated as the product of *DON* and *NIP* per site per year.

The biogeographic characteristics of the islands were expected to affect the presence and abundance of mammalian hosts on the islands. Thus, *the size of the island* (hectares) and *the minimum distance to the mainland* (meters) and *the minimum distance to neighbouring islands* (meters) from each study were identified in ArcGIS (version 10.8) (41).

### Statistical analyses

All statistical analyses were conducted in R version 4.4.3 (42). Linear mixed models (LMMs) were used to examine the effect of the study year on *rodent abundance, cervid dung abundance, DON* and *DIN*. A generalised linear mixed model (GLMM) was used to study the effect of study year on *NIP*, with island identity included as a random effect in the (G)LMMs. Predicted means and significant pairwise differences, based on Tukey’s test’s adjusted p-values, are reported.

#### Structural Equation Models (SEM)

To examine the direct and indirect relationships between the hosts (Year_t_) and the density of ticks and the pathogen in the following year spring (Year_t+1_, i.e. *DON*_*t+1*_, *NIP*_*t+1*_, *DIN*_*t+1*_), a structural equation modelling (SEM) approach was used. The structure of SEMs and explaining variables for each component model should be selected based on the biology of the system, (43) and the first step is to select the appropriate variables for the models (44). As both the presence and abundance of cervids may be relevant for the ticks, and because cervids were examined in spring and autumn, the most relevant cervid-related variables in the system were identified using single-variable regression models (45). Potential variables associated with DON_t+1_ were examined with linear regression using each of the following explanatory variables: (i) *cervid dung abundance in the previous spring (Year*_*t*_), (ii) *cervid dung abundance in the previous autumn (Year*_*t*_), (iii) *the presence (yes/no) of cervid dung in spring (Year*_*t*_) or (iv) *the presence (yes/no) of cervid dung in autumn (Year*_*t*_). The variables showing significant association with the response variable (*DON*_t+1_) (Table S2) were used in the SEM. The rest of the variables in the SEM were selected based on knowledge of the biology of the system and the study questions.

The SEM consisted of six component models to address the following relationships: **Model 1** (generalised linear model (GLM) with gamma error distribution) quantified *DIN*_*t+1*_ in association with *NIP*_*t+1*_ and *DON*_*t+1*_. **Model 2** (GLM with binomial error distribution and the number of tested nymphs as “weights” in the model to account for the differences in the examined nymphs) evaluated *NIP*_*t+1*_ in association with *rodent abundance* (pathogen-competent hosts) and *the abundance of cervid dung* in the preceding autumn (Year_t_), which coincided with when the nymphs (used in *DON*_*t+1*_, *NIP*_*t+1*_, *DIN*_*t+1*_) were questing and feeding as larvae. **Model 3** (GLM gamma) examined *DON*_t+1_ in association with *the presence of cervid dung* in the previous spring (Year_t_), *i*.*e*. the time when the females of the previous generation were questing for hosts. In addition, the potential effects of *the abundance of cervid dung* in the previous autumn (Year_t_) and *the rodent abundance* in the previous autumn (Year_t_) were examined in the same model. **Model 4** (linear model (LM)) examined *the rodent abundance* in autumn (Year_t_) in association with *the abundance of cervid dung* (at the same time point in autumn Year_t_) and *the presence of cervid dung* (in the spring Year_t_), *the island size, the minimum distance to the mainland*, and *the minimum distance to neighbouring island*. **Model 5** (LM) tested *the abundance of cervid dung* in autumn (Year_t_) in association with *the island size, the minimum distance to the mainland*, and *the minimum distance to neighbouring island*, and *presence of cervid dung* in the spring (Year_t_). **Model 6** (GLM with binomial error distribution) evaluated *the presence of cervid dung* in spring (Year_t_) in association with *the island size, the minimum distance to the mainland*, and *the minimum distance to neighbouring island. DON*_t+1_ and *DIN*_t+1_ was modelled with gamma distribution models, due to which a small number (0.001) was added to the observed values to avoid zero values. The SEM included rodent data from the autumns of 2019 and 2020, tick data from the springs of 2020 and 2021 and cervid data from springs 2019, 2020 and 2021 and autumns 2019 and 2020.

From the SEM, the significant paths in each model are reported, together with the R^2^, which describes the proportion of variation in the response variable explained by each model. The overall fit of the piecewise SEMs was evaluated by Fisher’s C statistic, which indicates whether there are any missing paths. Estimated coefficients, errors, and p-values of each model within the SEM are presented in Table S3. The SEM was fitted with the piecewiseSEM package version 2.3.0.1 (46). Each GLM and LM within the SEM was fitted with the stats package.

Rodent data from the autumn of 2017 and 2018 and tick data from spring 2018 and 2019 were not included in the SEM as the corresponding cervid data from 2017 and 2018 were not available. Therefore, the effect of *rodent abundance* in autumn (Year_t_) on *NIP*_*t+1*_, *DON*_*t+1*_, and *DIN*_*t+1*_ in the following year spring were examined using all available data (rodent data collected 2017-2020) using a GLM with gamma distribution (*DON*_*t+1*_, *DIN*_*t+1*_) and binomial distribution (*NIP*_*t+1*_).

## Results

### Rodent abundance across the islands

In total 1193 rodents were captured, of which 1145 (96 %) were bank voles (*Clethrionomys glareolus*), 25 were field voles (*Microtus agrestis*) and 23 were yellow-necked mice (*Apodemus flavicollis*). As the bank vole was much more abundant than the other rodents, and bank vole abundance (per 100 trap nights) was highly correlated with total rodent abundance (r=0.99, p<0.001) the inferences of the results based on rodent abundance are indistinguishable from the bank vole results. Rodent abundance varied greatly (0–140 individuals/100 trap nights) between the observations (site-year combinations), with overall mean of 32.5 individuals/100 trap nights. The abundance of rodents was relatively stable across the years, with the mean annual abundance being ca. 27 individuals/100 trap nights in 2017-2019, but significantly higher (mean 71 individuals/100 trap nights) in 2020 (Figure 2A, Table S3).

**Figure 2.**
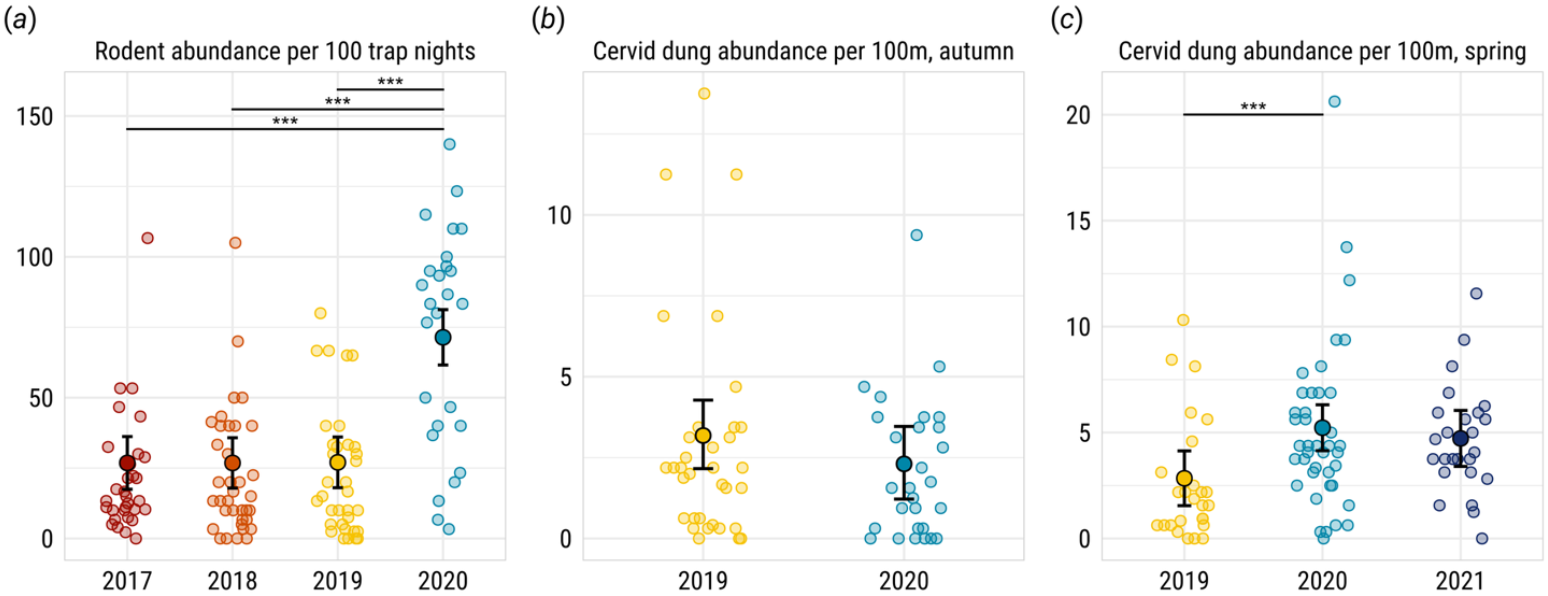
Observed (small filled circles) and model predicted values (larger filled circles) and their 95% confidence intervals (error bars) of A) abundance of rodents/100 trap nights, B) abundance of cervid dung piles in autumn and C) abundance of cervid dung piles in spring per year. Significant pair-wise comparisons (***p<0.001) between study years are identified with horizontal lines. The islands and sites that were examined were not identical between years.

### Cervid abundance across the islands

The cervid dung pile abundance was used as a proxy for the abundance of cervids (roe deer, white-tailed deer and moose). As the deer dung piles (combined roe deer and white-tailed deer) were much more abundant than moose dung piles, and deer dung abundance was highly correlated with cervid dung abundance (spring: r=0.94, p<0.001; autumn: r=0.95, p<0.001) the inferences of the results based on cervid dung abundance are dominated by the deer dung results.

The observed number of dung piles across the sampling sites varied from 0 to 20.6 (mean 4.25) dung piles/100m per site in spring and 0 to 13.8 (mean 2.57) dung piles/100m per site in autumn. The autumn dung count did not differ between the two study years (Figure 2B, Table S4), but the spring dung count was significantly higher in 2020 and marginally significantly higher in 2021 in comparison to 2019 (Figure 2C, Table S5).

### Total ticks and the density of nymphs (*DON*) in spring

In total 5942 nymphs, 493 adult females and 510 adult males were collected in the spring during 2018-2021. All nymphs were identified microscopically and a subset was confirmed using qPCR as *Ixodes ricinus*.

The density of nymphs (*DON*) in spring varied from 0 to 82.5/100 m^2^ per site per year, with a mean of 13.8/100 m^2^. In some islands, *DON* was relatively stable between sites and/or years whereas on some islands *DON* varied considerably between years. On one island (no. 5), *DON* was very high with an estimated mean of 53.5 nymphs/100 m^2^ and observed *DON* being 82.5/100 m^2^ in 2020 and 74.5/100 m^2^ in 2021. Consequently, the presented results are based on a dataset in which the observations (*n*=4) from this island were removed. However, inclusion of these observations did not change the main findings of the paper. The mean *DON* increased slightly over the years, being significantly higher in 2021 in comparison to the earlier study years (Figure 3A, Table S6).

**Figure 3.**
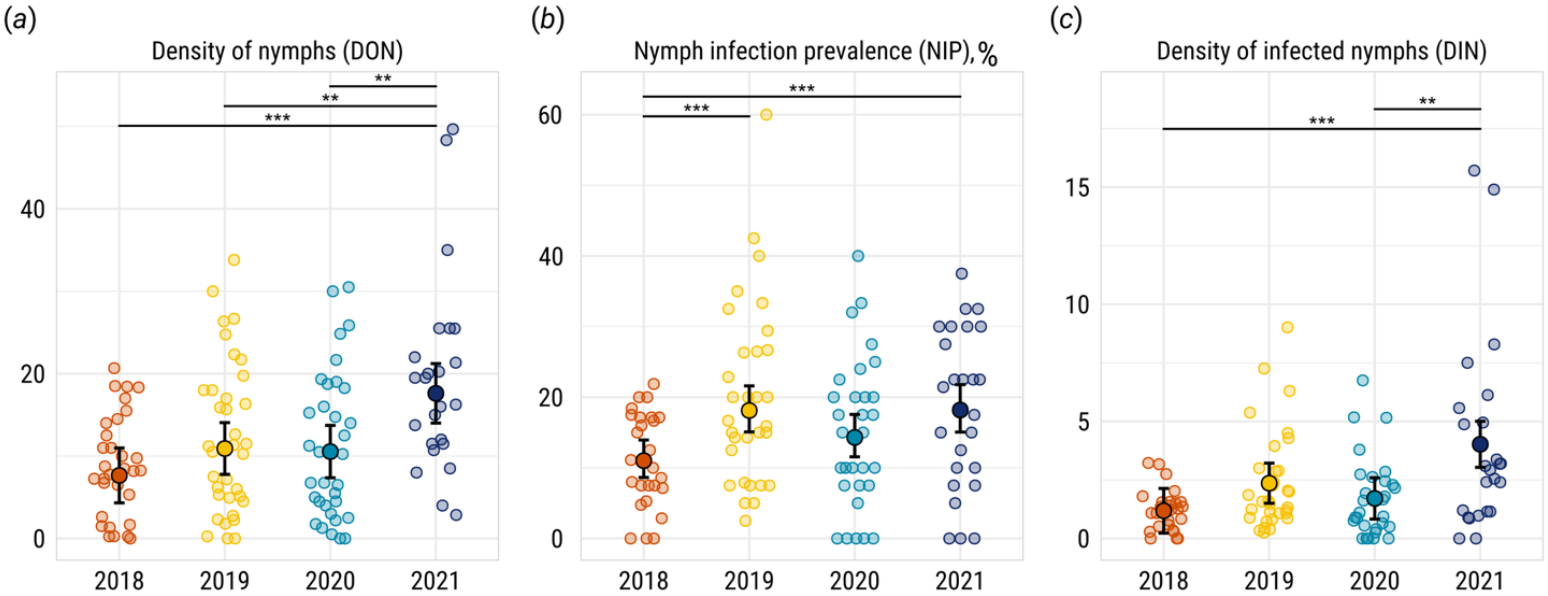
Observed (small filled circles) and model predicted values (larger filled circles) and their 95% confidence intervals (error bars) of A) density of nymphs (DON) per 100m dragging in the vegetation, B) *Borrelia afzelii* infection prevalence (%) in nymphs and C) density of *B. afzelii* -infected nymphs (DIN) per 100m in each study site per year. Significant pairwise comparisons (***p<0.001, **p<0.01) between study years are identified with horizontal lines. The islands and sites that were examined were not identical between years.

### *Borrelia afzelii* infection prevalence in nymphs (*NIP*)

In total 3742 nymphs were initially screened for *Borrelia burgdorferi* s. l. and 725 (19.4%) positive nymphs were further examined for *B. afzelii*. In total 597 nymphs were *B. afzelii* infected, which was 16% of all screened individuals and 82.4% of *B. burgdorferi* s. l. positive individuals, showing that *B. afzelii* is the dominating Lyme borreliosis-causing genospecies in the study system.

Nymphal infection prevalence (*NIP*) for *B. afzelii* was quantified for 113 observations (i.e., site-year combinations). The observed *NIP* varied between 0-60% per site per year, being on average 16.2%. *NIP* varied between years, being significantly higher in 2019 and 2021 in comparison to 2018 (Figure 3B, Table S7).

### Density of infected nymphs (*DIN*)

The density of *B. afzelii*-infected nymphs (*DIN*) per 100 m^2^ was estimated as the product of *DON* and *NIP* per site per year. *DIN* varied between 0-16.5 infected nymphs per 100 m^2^, with a mean of 2.4 infected nymphs/100 m^2^. *DIN* was the lowest in 2018 and the highest in 2021 (Figure 3C, Table S8).

### Associations between *Borrelia afzelii*, ticks, hosts and island characteristics

The density of *B. afzelii-*infected nymphs, *DIN*_*t+1*_, was positively associated with the density of nymphs (*DON*_*t+1*_) and the nymph *B. afzelii* infection prevalence (*NIP*_*t+1*_) at Year_t+1_. The variation in *DIN*_*t+1*_ was largely explained (R^2^=0.73) by the variation in *DON*_*t+1*_ and *NIP*_*t+1*_ (Figure 4, Figure 5A, 5B, Table S9).

**Figure 4.**
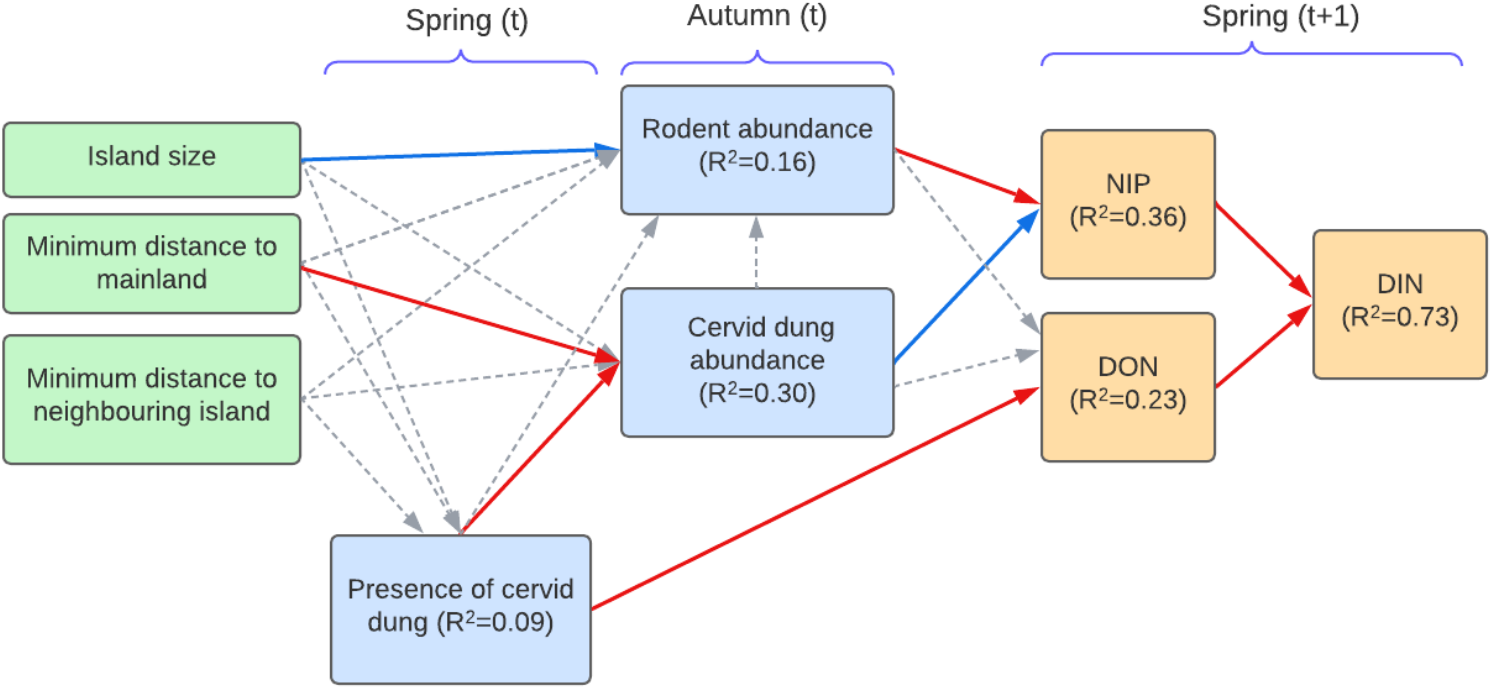
Path diagram of a piecewise Structural Equation Model (SEM) showing the effects of island characteristics (green boxes) on the mammalian hosts (blue boxes), and the effect of mammalian hosts on ticks (orange boxes), including nymph *B. afzelii* infection prevalence (NIP), the density of nymphs (DON), and the density of *B. afzelii* infected nymphs (DIN). Solid arrows represent statistically significant (p < 0.05) associations, with red indicating positive effects and blue indicating negative effects. Dashed grey arrows represent non-significant tested relationships. Fisher’s C = 22.852 with p-value = 0.410 on 22 degrees of freedom. The proportion of variation explained (R^2^) for each component model is given in the boxes of response variables.

**Figure 5.**
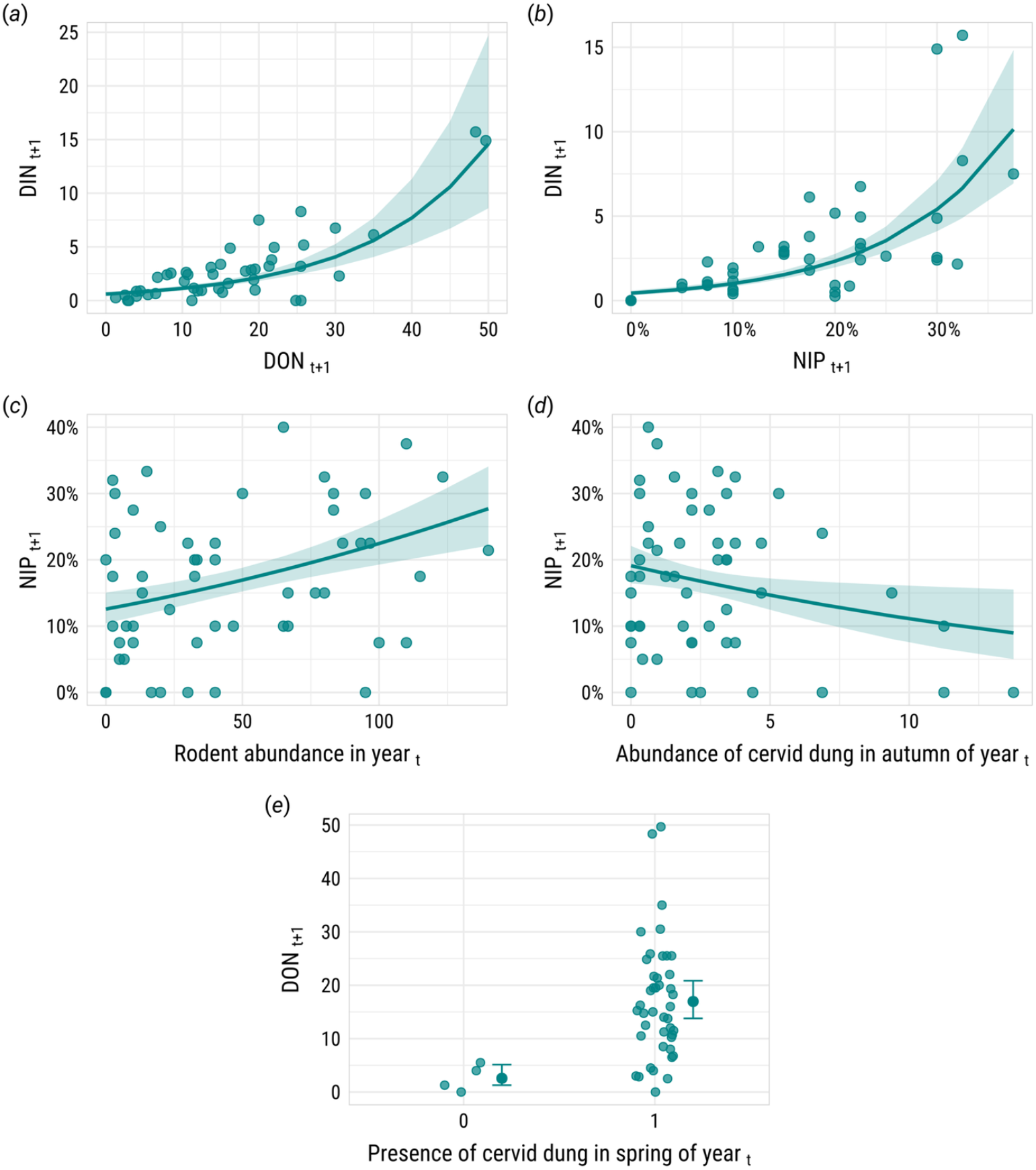
The observed (small filled circles) values and predicted (lines and the larger filled circles in +/-95% CI ribbons and error bars) significant associations identified in the compartmental models of the SEM. Model 1 examined the associations between density of *B. afzelii* infected nymphs in Year_t+1_ (DIN_t+1_) and simultaneously (A) density of nymphs (DON) and (B) nymph *B. afzelii* infection prevalence (NIP); model 2 examined the associations between NIP_t+1_ and (C) rodent abundance and (D) abundance of cervid dung in the previous autumn (Year_t_); model 3 tested the effect of the presence of cervid dung in previous spring (Year_t_) on (E) DON (Year_t+1_). In each model the other variables were set to the mean.

*NIP*_*t+1*_ was positively associated with *the rodent abundance* in the previous autumn (Year_t_) (Figure 4, Figure 5C) but negatively associated with *the abundance of cervid dung* in the previous autumn (Year_t_) (Figure 4, Figure 5D, Table S9). Due to the variation in the observed rodent abundances (from 0-140/100 trap nights), the positive effect of rodents resulted in an average increase in *NIP*_*t+1*_ from ca. 13 to 27% (Figure 5C). *Cervid dung abundance* in autumn, however, varied from 0-13.75 dung piles/100 m, which decreased *NIP*_*t+1*_ from ca. 19% to 9% (Figure 5D). The abundance of rodents and cervid dung together explained 36% of the variation in *NIP*_*t+1*_ (R^2^=0.36).

*DON*_*t+1*_ was positively associated with the presence of cervid dung on the site in the previous spring (Year_t_), but not *the abundance of rodents* or *cervid dung abundance* in the previous autumn (Figure 4, Figure 5E). *The presence of cervid dung* increased *DON*_*t+1*_ from 2 nymphs/100 m^2^ (no cervid dung observed on the site) to ca. 17 nymphs/100 m^2^ (cervid dung observed on the site) (Figure 5E). The model explained 23% of the variation in *DON*_*t+1*_ (Figure 4).

*Rodent abundance* in autumn (Year_t_) was not associated with *the presence of cervid dung* in the spring or *abundance of cervid dung* in autumn (Year_t_), suggesting that the two hosts are not associated with each other (Figure 4, Table S9). Rodent abundance though was negatively associated with island size, with rodent abundance being higher on smaller islands (Table S9, Figure S1). However, rodent abundance was not associated with isolation (*distance to mainland or neighbouring islands*) (Table S9). Together t*he presence and abundance of cervid dung* and island characteristics accounted for 16% of the variation in *rodent abundance* (Figure 4).

*The presence of cervid dung* on a site was not associated with any of the examined island characteristics, and only 9% of the variation was explained by the island characteristics (Figure 4). *The abundance of cervid dung* in autumn, however, was positively associated with *the minimum distance to the mainland* and *the presence of cervids* on the site in the previous spring (Figure 4, Table S9, Figure S2), and 30% of the variation in *cervid dung abundance* was explained by the model.

The SEM results were further supported when all available rodent data (2017-2020) were used. There was no association between *rodent abundance* in autumn (Year_t_) and *DON* in the spring of the following year (*DON*_t+1_), whereas a significant positive association was detected between *the rodent abundance* and *NIP*_*t+1*_ and *DIN*_*t+1*_ (Table S10, Figure S3).

## Discussion

The aim of this study was to quantify the role of rodents and cervids, the key reproductive hosts for the pathogen and the tick, respectively, in determining the hazard of Lyme borreliosis in boreal Fennoscandia, where rodent populations show dramatic density variations. Structural Equation Modeling (SEM) were used to capture the complexity of this disease system where multiple factors interact through both direct and indirect pathways and determine LB hazard. With our extensive field study across 41 islands, in which the abundance of rodents, cervids and ticks, and the prevalence of pathogen in ticks varied greatly, we show that the prevalence and density of infected nymphs are driven by the rodents that amplify the pathogen in the ticks. However, cervids have a dual role, both amplifying the density of nymphs, but also diluting the prevalence of infected nymphs, supporting our hypothesis of the roles of different types of hosts in the disease system. However, contrasting our hypothesis on the role of rodents in supporting ticks, the density of nymphs was not associated with the preceding year’s rodent abundance, suggesting that in this system rodents contribute primarily to pathogen transmission, rather than tick maintenance. Importantly though, very low abundance or virtual absence of rodents, did not cause the pathogen to fade out from the following nymph tick cohort, despite the decrease in the infection prevalence at lower rodent densities. Moreover, within the observed range of cervid abundance, dilution of the pathogen was observed, with a negative relationship between autumn cervid abundance and nymph infection prevalence the following spring. Meanwhile, the presence of cervids was the key determinant of the tick abundance in the system. Our data highlight, and help to unpick, the complex interactions that affect LB hazard in the northernmost zone of the tick distribution. In particular, our results suggest that despite the extensive between-year and between-site variations in host densities, the population dynamics of these key host groups do not interrupt the transmission and maintenance of the pathogen in this system.

### Rodents drive the pathogen circulation

Our data showed a positive association between rodent abundance and the prevalence of *B. afzelii* in nymphs the following year. This finding mirrors several earlier studies showing that higher rodent densities are associated with higher prevalence of rodent-associated tick-borne pathogens in nymphs (9,10,18,19). However, our study system is likely to differ from most earlier studies as the rodents in our study system exhibit very strong abundance variations. Most importantly, no rodents were caught in the autumn on several occasions, suggesting that the rodent population was absent or at very low densities over summer, the breeding season of rodents and an important feeding period for larvae. The absence of the key hosts of a pathogen is expected to prevent the pathogen’s transmission from the hosts to ticks (2,47). However, the absence of rodents in the autumn trapping did not result in the absence of the pathogen in nymphs the following summer. This is in line with an earlier experimental study showing that removal of rodents over their breeding season did not decrease pathogen prevalence in the ticks on the islands (37). This may be because some uncaptured rodents or other small mammals that support the transmission of the pathogen were present in the islands. Indeed, red squirrels and shrews, which are feeding hosts for immature ticks and competent hosts for *B. afzelii* (48–51), have been detected in blood meal analyses of nymphs in some of the islands (52), but their abundances were not estimated in this study. Alternatively, at low rodent densities ticks may aggregate on the few small mammal individuals that are present, potentially facilitating the pathogen’s transmission (50, 53). Although our findings suggest that low rodent density during the larval tick activity period is not sufficient to interrupt the natural circulation of the pathogen, lower rodent abundance was associated with lower infection prevalence, suggesting that the low phase of the rodent cycles may hinder pathogen transmission, reducing its overall prevalence in the system (47).

While low rodent density was expected to be a critical factor for the persistence of the pathogen in the system, higher rodent densities are generally expected to facilitate the pathogen’s transmission (47,54). However, it has also been suggested that very high rodent abundances may hinder pathogen transmission, as tick infestation load per rodent may decrease, and a higher proportion of larvae may feed on uninfected rodents, causing decrease in the subsequent year’s nymphal infection prevalence (54). Further studies are needed to quantify the potential nonlinear associations between rodent abundance and pathogen circulation in cyclic rodent populations.

### The effect of rodents and cervids on the vector population

Rodents are regarded as important hosts for larval ticks, and in several studies, DON has seen to be positively associated with rodent density in the previous year (8,9). However, we found no association between the abundance of rodents and DON in the following year. One reason could be that alternative hosts may support larval feeding, thus decreasing the relative importance of rodents. Indeed, another study in the islands showed that while rodents provide more than 50% of all larval blood meals, ca. 12 % of larvae had fed on deer and ca. 10-12% had fed on squirrels and shrews, respectively (52). Cervids have been widely identified as important hosts for immature ticks in other study systems (6,25). The importance of cervids as the hosts of larval ticks was further supported by the observed decrease of NIP with the increase of cervids in our study. In other words, the more cervids the bigger the proportion of larval ticks that feed on the cervids rather than on rodents, and since cervids are non-competent hosts for the pathogen, the lower the infection prevalence become. However, we did not detect an association between DON and cervid dung abundance in the previous year, suggesting that although cervids are important hosts for larval ticks, DON is primarily dependent on other drivers than the success of larval ticks.

Our findings indicate that mere presence of cervids in spring and not their abundance was associated with higher nymph density, in line with earlier studies showing that the presence of deer is a strong determinant on the density of nymphs (19,27). Indeed, cervid densities varied drastically between the islands, where absence of cervid dung in spring on some islands was followed by low DON the following year. In such cases, it seems likely the low DON result from female ticks being unable to feed and reproduce in the absence of cervids, rather than from a lack of hosts for larval feeding. Consequently, the density of nymphs in our study system appears to be primarily dependent on the availability of sufficient hosts supporting tick reproduction, rather than on the hosts that support the development of immature ticks.

### Dilution of pathogen prevalence in ticks caused by cervids

It has been suggested that a high abundance of hosts that are not competent to transmit the pathogen may decrease the proportion of ticks feeding on competent hosts for pathogen transmission, leading to a decrease of infection prevalence in the following life stages, causing the dilution effect (6,7,55). We observed a significant negative association between the abundance of cervids and *B. afzelii* infection prevalence in nymphs in the following year, suggesting that cervids were diverting larval ticks from feeding on pathogen competent hosts and, thus, causing a dilution effect. Theoretical studies have shown that such a dilution effect, caused by high density of hosts that are not competent transmission hosts for the pathogen, may cause the pathogen to fade out from the ecosystem (1,22,47). In our study, *B. afzelii* prevalence indeed decreased and was even zero in two islands with the highest cervid dung counts. Interestingly, these two islands are relatively large and next to each other, and together with a few neighbouring islands they likely support their own cervid population. However, in both islands, DON was relatively low (<3 nymphs/100 m2) in the occasions no pathogen was detected. As such, the estimated low infection prevalence may either be a result of the low number of tested nymphs (i.e., a sampling effect), or the low density of ticks that hinder the transmission of the pathogen (i.e., a limitation on transmission). Nevertheless, there was a clear overall decrease in NIP in association with cervid dung abundance. Mathematical modelling of the system would be required to further explore the cervid densities needed for the dilution effect to be strong enough that the pathogen is eliminated from the system.

### LB hazard and the role of rodents and cervids

Density of infected nymphs (DIN) is the most relevant measure for the hazard of LB (10) and DIN is the product of the density of nymphs (DON) and nymph infection prevalence (NIP). In our study islands, the rodent abundance was relatively high (average 32.5/100 trap nights) and was positively associated with NIP, highlighting the increasing effect of rodents on LB hazard. However, rodents cannot support the abundant tick population, for which the cervids are required. As Gandy and coworkers (6) proposed, LB hazard (=DIN) is the highest in environment with high rodent density (support high NIP) and high deer density (support high DON). However, in their study high deer density decreased rodent density due to cascading effects through vegetation, suggesting that the high abundances of both rodents and deer are unlikely (6). Thus, DIN may be limited either by lower density of cervids (affect through DON) or lower density of rodents (affect through NIP) (6). In our study, however, rodent abundance was not associated with the abundance of cervids, potentially because bank voles were dominating rodent community and they are generalist that use diverse food items. Therefore, boreal forests may support both high cervid density but also high rodent density, supporting high DON and high NIP, which translate into high LB hazard.

The role of cervids in LB hazard is more complex, as high levels of cervids results in vector amplification but pathogen dilution. This dual effect means the overall impact of cervids on disease hazard may be nuanced, with countervailing forces sometimes neutralising each other (7,56). In our study, cervid presence determined tick abundance, whereas pathogen dilution was dependent on the cervid abundance. Therefore, in our study system the counteracting effects of cervids are unlikely to neutralise each other, as even the relatively low abundance of cervids support ticks, whereas the decrease in NIP was clear only in cervid dung abundances higher than average (>5 dung piles per 100m^2^). Consequently, the overall effect of cervids likely increased, rather than decreased, LB hazard in our study system.

## Conclusions

Harnessing the natural variation in rodent and cervid densities across our boreal island system, we show that the hazard of rodent associated tick-borne pathogens is determined dynamically by the availability of different types of hosts that provide the blood meals for ticks and transmission opportunities for the pathogen, respectively. In particular, the density variations of rodents translated into variation of LB hazard, but the periodic lack of rodents was not sufficient to result in elimination of the pathogen. Moreover, the overall effect of cervids is likely to increase the hazard, despite the observed dilution effect on NIP, although the exact nature of the interplay needs further quantification. These inferences were enabled by the high degrees of natural variation in deer and rodent densities across our island system. To achieve similar variations in mainland forest systems would require extensive spatial (large area) and temporal (over long time period) sampling, increasing the likelihood that other biotic and abiotic conditions would covary with the hosts.

Applying our SEM framework to the natural variation in host, tick and pathogen densities across island ecosystems provided a mechanistic understanding of the complex host–tick–pathogen system, enabling identification of the roles of different types of hosts in driving LB hazard in boreal forests.

## Acknowledgements

We thank Tanja Hirvonen, Katariina Tiainen for their help in the field, Sonja Knuutila and Milla Rajala for their help in the lab. We thank Camilla Sundbäck, Mikael Sundbäck and Johan Sundbäck for the help with recruiting islands to the study, transportation and help in the field.

## Funding

We want to thank all the landowners for the access to their land to carry out the study. This research was supported by the funding from the Research Council of Finland (project numbers 335651, 329326 and 354988 to ERK and 324605 to TM). NK was funded by the University of Jyväskylä Graduate School, with additional support by the Societas pro Fauna et Flora Fennica.

## Author Contributions

ERK, AF, TM conceived the ideas and designed methodology; YXGW, NK, IB, SS, ET, SV, EW, TM and ERK collected the data; NK, SS, JK, SV and EW carried out laboratory analyses, YXGW and ERK analysed the data; YXGW, NK, AF and ERK led the writing of the manuscript. All authors contributed critically to the drafts and gave final approval for publication.

## Notes

### Competing Interest Statement

The authors have declared no competing interest.

### Summary of Updates

Author name was incorrectly spelled, but is now fixed.

