## Supplementary Information for "Contrasting roles of host species in tick-pathogen interactions and their influence on Lyme borreliosis hazard across boreal island communities"

**Corresponding to:** eva.r.kallio @jyu.fi

**Affiliations:**

1. Department of Biological and Environmental Science, University of Jyväskylä, Finland
2. Institute for Nuclear Research of the NAS of Ukraine, Ukraine
3. Institute of Infection, Veterinary & Ecological Sciences, University of Liverpool, UK

**ORCID:**

Yingying X.G. Wang: 0000-0003-3066-197X
Nosheen Kiran: 0009-0001-5125-2932
Ilze Brila: 0000-0002-1518-1380
Saana Sipari: 0000-0001-8846-1538
Saija Vuorenmaa: 0009-0001-9247-8620
Andy Fenton: 0000-0002-7676-917X
Tapio Mappes: 0000-0002-5936-7355
Eva R. Kallio: 0000-0003-2991-612X

### Supplementary methods

#### Study area

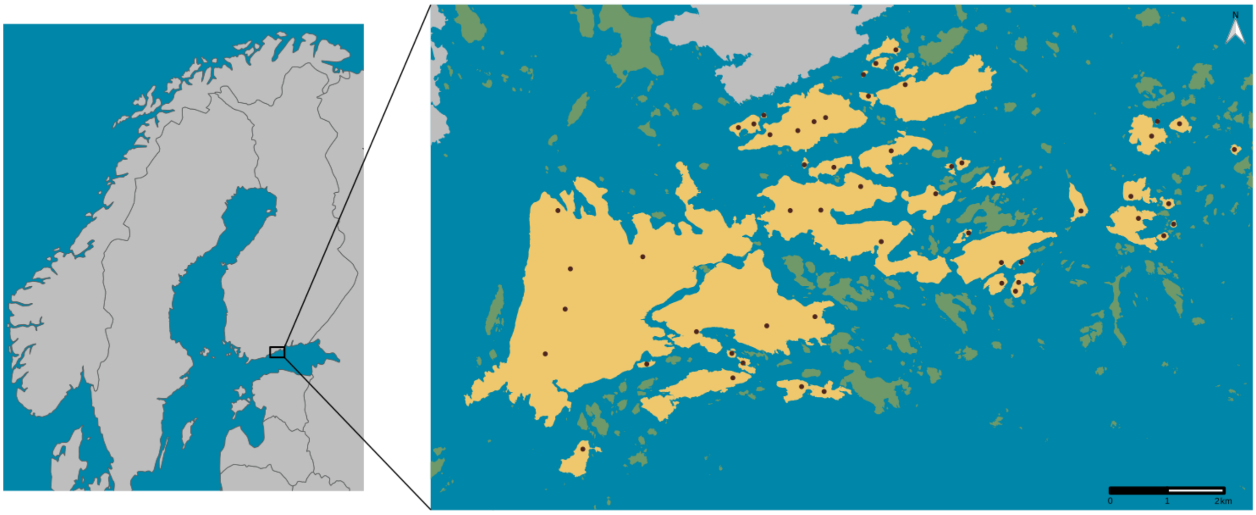

Map S1. Study islands (yellow) and sites (red dots) in Porvoo archipelago, including the mainland (grey) and islands that were not included in the study (green).

#### DNA extraction

DNA was extracted using ammonium hydroxide method, in which ethanol stored nymphs were air dried and transferred to 200 µl of 1.25 % NH4OH (ammonium hydroxide) solution. The sample solution was preheated (3 minutes at 100 °C), crushed with a 5 mm steel bead using a lyser (2x1min at 26 Hz) (Qiagen TissueLyser, USA), and incubated at 100 °C for 20 minutes. The tubes were cooled at room temperature and centrifuged briefly before being opened and incubated at 100°C to allow the ammonia to evaporate until desired extraction volume (50% of starting volume).

#### Pathogen detection

The PCR mix for all qPCRs consisted of 7.5 µl Itaq universal Probes Supermix (Bio-rad, USA), 0.75 µl of primers (10 µM), 0.375 µl of probe (10 µM), 2 µl of Bovine albumin serum (BSA) (5 mg/ml) and 3.5 µl of nymphal DNA sample making up to a final volume of 15 µl with sterile molecular grade water. Cycling conditions in the BioRad CFX96 instrument were 5 min denaturation at 95 °C, followed by 50 cycles for *B. burgdorferi* s.l. and 45 cycles for *B. afzelii* with denaturation at 95 °C for 10 sec and annealing at 60 °C for 1 min. In addition to the samples and negative extraction controls, each 96 well plate included one negative control with only dH2O and 1-2 positive controls. If any negative extraction controls showed a positive signal in the qPCR, five samples before and after the potentially contaminated negative controls were repeated. After the repeat, no contaminations were detected.

**Tick species identification with qPCR**

A subset of 350 nymphs was identified for tick species using a species-specific duplex real-time quantitative PCR (qPCR) assay (Laaksonen et al. 2018). The primers and tick species-specific probes known as Ipe-I2-P4 for *Ixodes persulcatus* and Iri-I2-P4 for *I. ricinus* are provided in Table S1. The qPCR mix contained 2.1 µl molecular biology grade water (Sigma) 0.2 µl of each primer (10 µM), 0.1 µL of Ipe-I2-P4 probe (10 µM), 0.15 µL of Iri-I2-P4 probe (10 µM), 1.25 µL of BSA (5 mg/mL) and 5 µL of ITaq universal Probes Supermix (Bio-rad, USA). PCR amplification was carried out with an initial 5 min denaturation at 95 °C followed by 40 cycles of denaturation at 95 °C for 10 sec and annealing at 60 °C for 30 sec.

**Supplementary results**

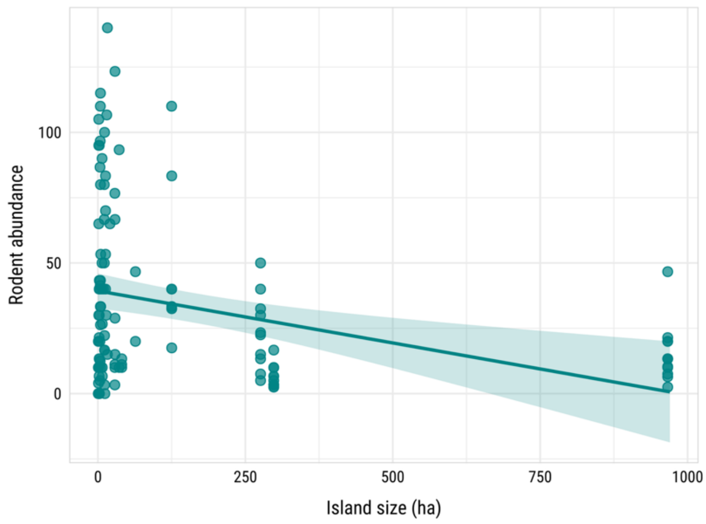

Figure S1. The observed (filled circles) values and predicted (lines with +/- 95% CI ribbons) significant associations identified in Model 4 of the SEM. Model 4 examined the associations between rodent abundance and the size of the island (in hectares).

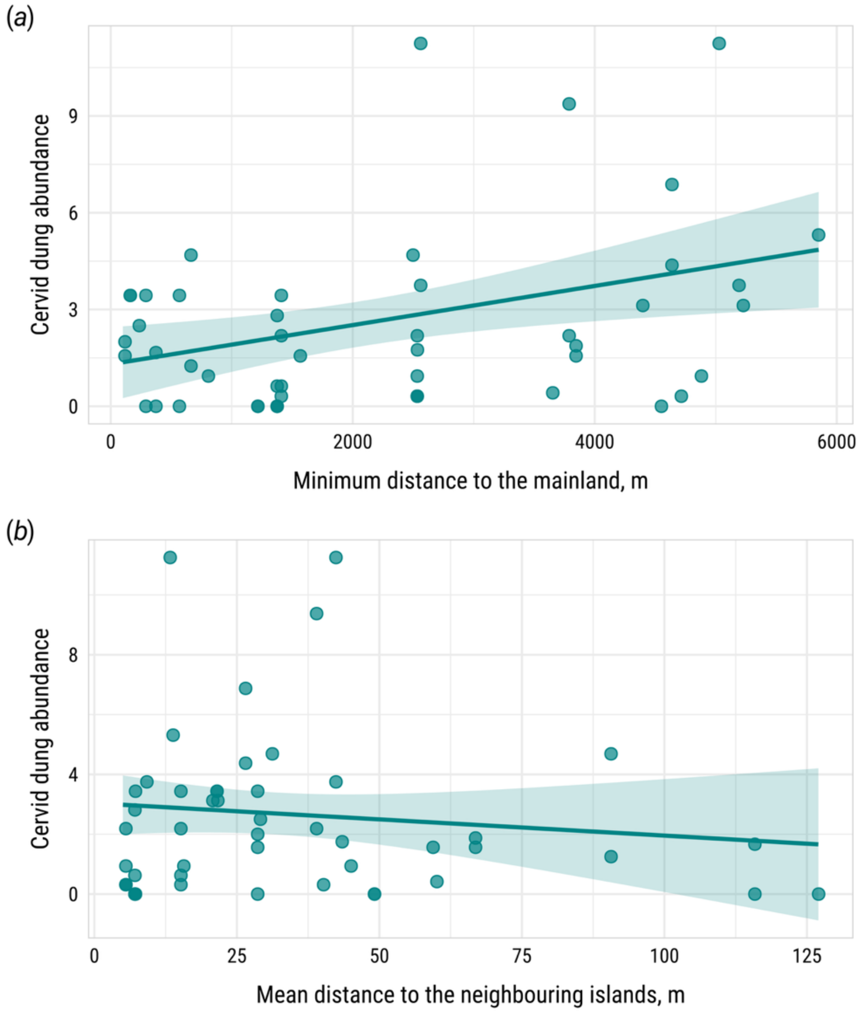


Figure S2. The observed (filled circles) values and predicted (lines with +/- 95% CI ribbons) significant associations identified in Model 5 of the SEM. Model 5 showed significant associations between the abundance of cervid dung in autumn and A) the distance from the island to the mainland (in meters) and B) the mean distance to neighbouring islands within 500 m radius.

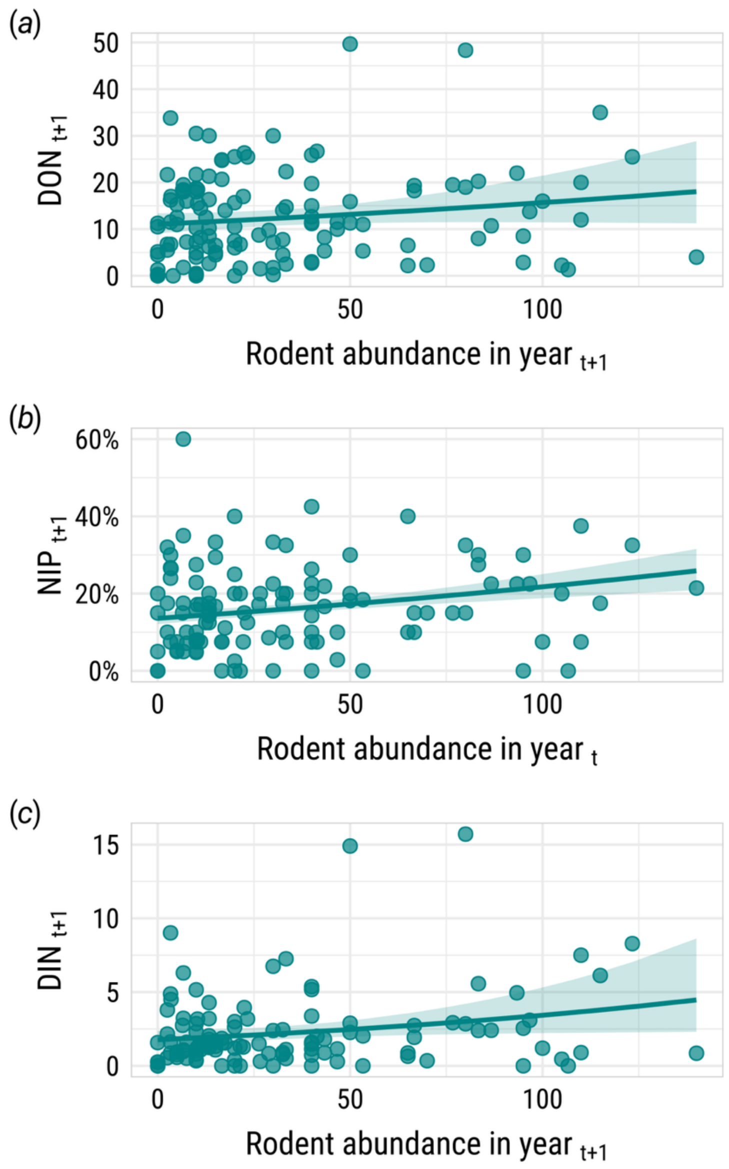


Figure S3. The observed (filled circles) values and predicted (lines with +/- 95% CI ribbons) significant associations between rodent abundance in autumn ($Year_t$) and the following year's (A) DON, (B) NIP and (C) DIN, based on rodent data collected in 2017-2020.
